# *O*-fucosylation of thrombospondin-like repeats is required for processing of MIC2 and for efficient host cell invasion by Toxoplasma gondii tachyzoites

**DOI:** 10.1101/382515

**Authors:** Giulia Bandini, Deborah R. Leon, Carolin M. Hoppe, Yue Zhang, Carolina Agop-Nersesian, Melanie J. Shears, Lara K. Mahal, Françoise H. Routier, Catherine E. Costello, John Samuelson

## Abstract

*Toxoplasma gondii* is an intracellular parasite that causes disseminated infections which can lead to neurological damage in fetuses and immunocompromised individuals. Microneme protein 2 (MIC2)^2^, a member of the thrombospondin-related anonymous protein (TRAP) family, is a secreted protein important for motility, host cell attachment, invasion, and egress. MIC2 contains six thrombospondin type I repeats (TSRs) that are modified by C-mannose and O-fucose in *Plasmodium* spp. and mammals.

Here we used mass spectrometry to show that the four TSRs in *T. gondii* MIC2 with protein *O*-fucosyltransferase 2 (POFUT2) acceptor sites are modified by a dHexHex disaccharide, while Trp residues within three TSRs are also modified with *C*-mannose. Disruption of genes encoding either *pofut2* or nucleotide sugar transporter 2 (*nst2*), the putative GDP-fucose transporter, results in loss of MIC2 O-fucosylation, as detected by an antibody against the GlcFuc disaccharide, and markedly reduced cellular levels of MIC2. Furthermore, in 10-15% of the *Δpofut2* or *Δnst2* vacuoles, MIC2 accumulates earlier in the secretory pathway rather than localizing to micronemes. Dissemination of tachyzoites in human foreskin fibroblasts is reduced in these knockouts, which both show defects in attachment to and invasion of host cells comparable to the phenotype observed in the Βmic2.

These results, which show *O*-fucosylation of TSRs is required for efficient processing of MIC2 and for normal parasite invasion, are consistent with the recent demonstration that *P. falciparum Δpofut2* has decreased virulence and support a conserved role for this glycosylation pathway in quality control of TSR-containing proteins in eukaryotes.

## Introduction

*Toxoplasma gondii* is a eukaryotic parasite that belongs to the Apicomplexa phylum, together with other medically relevant parasites such as *Plasmodium* spp. and *Cryptosporidium parvum. T. gondii*, which has the ability to invade all warm blooded animals, causes transient, disseminated infections that can lead to neurological damage in immunocompromised individuals and developmental defects in fetuses (1, 2). *T. gondii* replicates asexually in all of its hosts, with the exception of felids (cats), where the parasite also undergoes a sexual cycle, which concludes in production and shedding of infectious oocysts in the feces (3).

Because *T. gondii* is an obligate, intracellular pathogen, its virulence is dependent on the ability to invade host cells. The parasite has specialized secretory organelles, which localize at the apical end of asexual stages and whose content is sequentially secreted during the invasion process (4). Secretion of microneme proteins is dependent upon elevation of parasite calcium levels and is the first event upon parasite attachment (4, 5).

Microneme protein 2 (MIC2) is a member of the thrombospondin-related anonymous protein (TRAP) family (6). MIC2 is a type 1 membrane protein composed of an A/I von Willebrand domain, six thrombospondin-like repeats, and a short cytoplasmic tail (7). The A/I domain has been shown to be involved in recognition of host cells, possibly through binding to ICAM-1, heparin, and sulfated glycosaminoglycans (8-10). MIC2 forms a hexameric complex with the MIC2-associated protein (M2AP) in the endoplasmic reticulum; together they traffic to the micronemes and, upon secretion, to the apical cell surface (8, 11). As host cell penetration concludes, the MIC2/M2AP complex relocates to the posterior end of the parasite and is released into the environment by cleavage of the C-terminal transmembrane helix that is catalyzed by the MPP1 protease (7, 12).

Although knockout of the *mic2* gene strongly affects motility and invasion, MIC2 is not essential for parasite viability. Tachyzoites lacking this protein still glide, attach to, and invade fibroblasts, albeit inefficiently (13). Numerous studies have shown that correct trafficking of the M2AP/MIC2 complex to the micronemes is important for its function. Knockout of *m2ap* results in inefficient localization of MIC2 to the micronemes and defects in the parasite’s ability to glide and invade (14). Cleavage-resistant mutations of *m2ap*, which affect protein maturation and consequently localization of the M2AP/MIC2 complex, also result in inefficient gliding and attachment (15, 16).

Thromobospondin-like repeats, which are each about 60 amino acids long, are present in many extracellular proteins. TSRs have a conserved structure characterized by a tryptophan ladder composed of stacked Trp and arginine residues and a set of conserved cysteines that are involved in disulfide bridges (17). In metazoans, TSRs are glycosylated via two pathways: *O-*fucosylation by action of the protein *O*-fucosyltransferase 2 (POFUT2) and C-mannosylation by the C-mannosyltransferase DPY-19 (18, 19). C-Man is transferred via an unusual carbon-carbon bond to the Trp on a WXXWXXC^1^ sequence, where C^1^ is the first conserved cysteine in the TSR (20). *P. falciparum* and *T. gondii* each encode for a DPY-19 orthologue that can transfer *C*-Man to the Trp residues in the conserved motif, both *in vitro* and *in vivo* (21).

POFUT2 transfers a-fucose to the hydroxyl group on Ser/Thr residues in the consensus sequence C^1^X_2-3_(S/T)C^2^X_2_G, where C^1^ and C^2^ are the first two conserved cysteines and X is any amino acid (22). In metazoans, fucose is transferred on already folded TSR domains (23) and can be elongated by a ß1,3-glucose by action of the glucosyltransferase B3GLCT (24). Mass spectrometry analyses of *P. falciparum* TRAP and circumsporozoite protein (CSP) indicate that a hexose (Hex) residue elongates the fucose on Apicomplexa TSRs; this sugar may be glucose, but has not yet been unequivocally identified (21, 25).

TSRs glycosylation is carried out in the endoplasmic reticulum (22, 26) and, in studies performed in mammalian cell and *C. elegans*, defects in both *C*-mannosylation and *O*-fucosylation have been shown to affect protein stabilization and secretion, albeit with protein-to-protein variability (19, 23, 27-31). A knockout of *pofut2* is embryonically lethal in mice (31). The human recessive disorder Peter Plus syndrome is caused by mutations in the B3GLCT glycosyltransferase (23). Recently, a knockout of *P. falciparum pofut2* showed O-fucosylation of TSRs is required for efficient mosquito infection by ookinetes and for hepatocyte invasion by sporozoites (32). These phenotypes are due to defective folding/stabilization of the parasite TSR-containing proteins, such as TRAP and CSP, consistent with the observations made in metazoans and a potential role for *O*-fucosylation in efficient folding and/or stabilization of TSRs (23, 32).

In this paper we use mass spectrometry to show that four of six TSRs of MIC2 are modified by O-fucose, and that in addition to TSR5 (shown Hoppe *et al.*), two other TSRs are modified with C-Man. We also identify the putative POFUT2 and GDP-fucose transporter (NST2) and show that knockout of either *pofut2* or *nst2* results in loss of O-fucosylation of MIC2, defects in the protein stability and localization, and decreases the ability of parasites to invade host cells.

## Results

### Four TSRs of MIC2 are modified by a dHex with or without Hex

Glycosylation on tachyzoite secreted proteins was analyzed as described in (25). Briefly, secretion was induced with 1% ethanol (7), and the peptides resulting from reduction, alkylation, and trypsin digestion of the secreted protein fraction were analyzed by nanoLC-MS/MS using a Thermo Q Exactive quadrupole Orbitrap MS with low energy CID. Following this discovery run, selected peptide species containing the *O*-fucosylation motif were further analyzed by higher collision energy dissociation (HCD) fragmentation and by electron transfer dissociation (ETD), which allows fragmentation of the peptide backbone while preserving post-translational modifications (33).

The initial analysis identified numerous peptides from MIC2 that cover about 66% of the secreted sequence (7) (Fig. S1). *In silico* mining of the *T. gondii* predicted proteome indicated the possibility for detection of three additional TSR-containing proteins encoding the POFUT2 consensus motif: MIC14, TGGT1_209060, and TGGT1_223480 (Table S1). However, we did not observe any peptides from these three *T. gondii* proteins.

MIC2 has six contiguous TSRs. All six contain *C*-mannosylation sites, whereas only four (TSR1, 3, 4, and 5) have the consensus O-fucosylation motif (C^1^X_2-3_(S/T)C^2^X_2_G) (Fig. 1A). The targeted HCD and ETD experiments allowed localization of a dHexHex disaccharide on the predicted POFUT2 acceptor residues on MIC2 TSR1 and TSR4 and a dHex +/-Hex on TSR5 (25). TSR3 is also modified with a dHexHex, although the modification site could not be assigned (Table 1, Figs. 1, 2, 3 and S3). TSR6 lacks a consensus site for POFUT2 and is not modified by C-Man; TSR2 lacks an O-fucosylation consensus site and was not sampled. TSR3 and TSR4, in addition to the previously described TSR5 (25), are modified with C-Man. Lastly, while MIC2 contains four potential *N*-glycosylation sites, we observed peptides containing unmodified Asn at residues 463, 470, and 680, suggesting that these N-glycan sites are unoccupied (Fig. S1). The fourth N-glycosylation site was not observed.

**Table 1:**
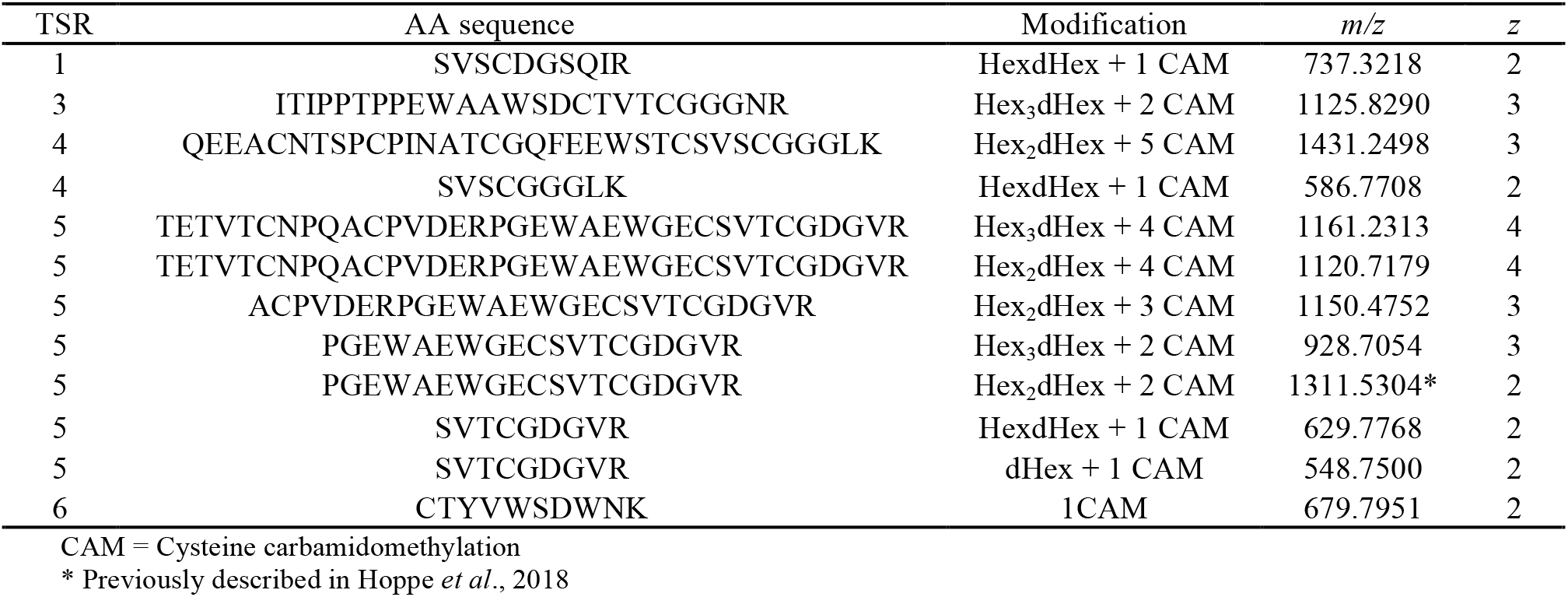
Peptides and glycopeptides from the TSRs of MIC2 detected in this study.

**Figure 1:**
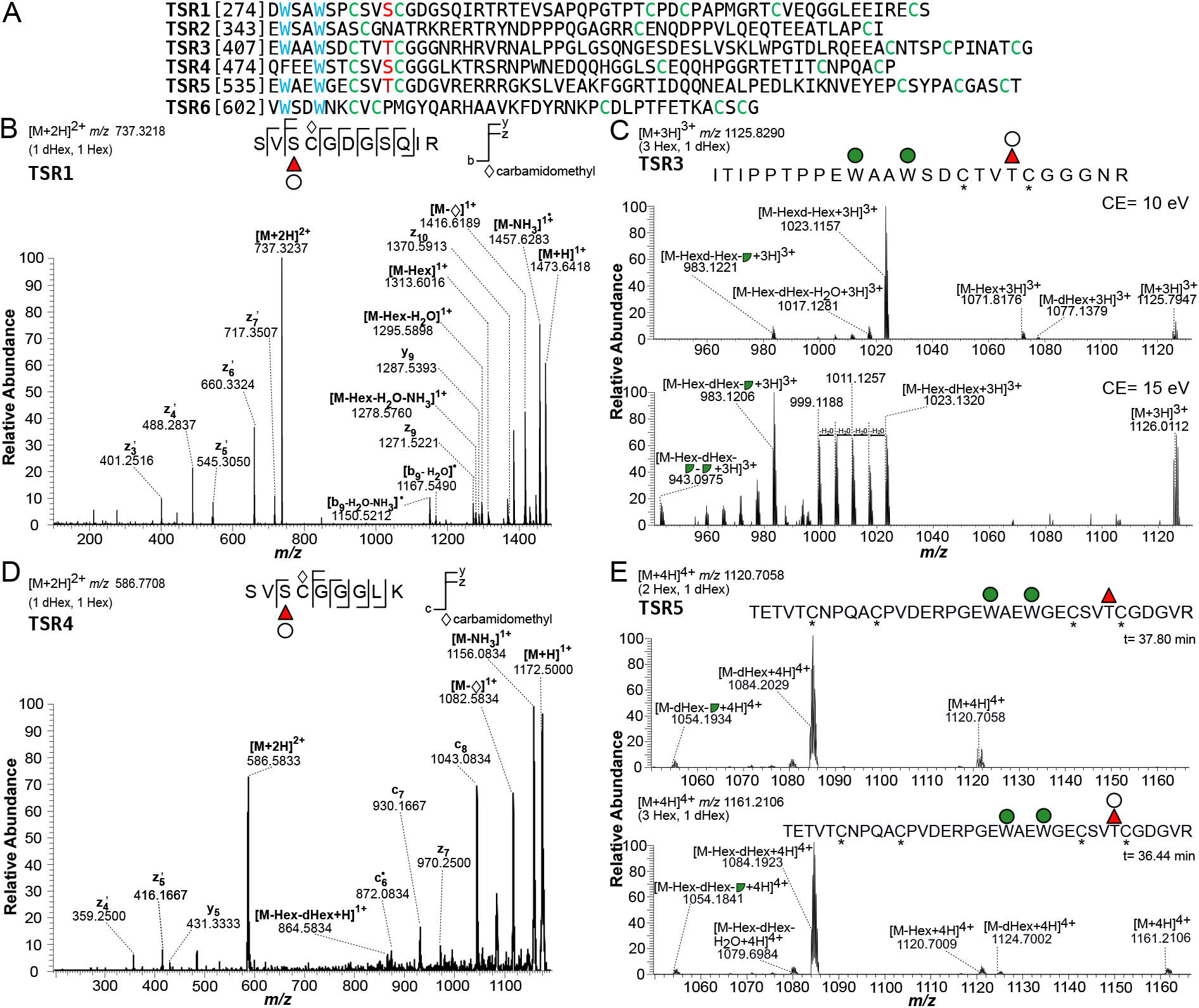
MIC2 TSR1, 3, 4, and 5 are modified by dHexHex and TSR3, 4, and 5 are also C-mannosylated. A. Amino acid sequences of the 6 TSRs domains of MIC2. Cys resides are labeled green, the O-fucosylation acceptor Ser/Thr is marked in red. C-mannosylation sites are shown in blue. B. ETD of a semi-tryptic peptide from TSR1. The presence of the three product ions z9, z10, and y10 +dHexHex identifies Ser285, the predicted POFUT2 acceptor as the glycosylated residue. *C*. HCD fragmentation of a TSR3 semi-tryptic peptide is consistent with a dHexHex disaccharide on plus two C-Mannoses. Collision energy of 10eV (top panel) shows ion species corresponding to the precursor minus a Hex, a dHexHex, or minus dHexHex and 120.04 Da (broken green circle). Increasing the collision energy to 15 eV (bottom panel) allows detection of the precursor minus Hex, dHexHex, and two Hex -120.04 Da fragments. *D*. IonTrap ETD of a semi-tryptic peptide from TSR4 identified the z7 product ion carrying both Hex and dHex, indicating Ser485, the predicted POFUT2 acceptor site, as the residue modified by a dHexHex disaccharide *E*. 10 eV HCD fragmentation of the Hex2dHex glycoform, *m/z* 1120.7058 (top panel) and the Hex3dHex ion species, *m/z* 1161.2106 (bottom panel) is shown. The most abundant ion, corresponding to the precursor minus the *O*-linked sugar, was the same for both species, consistent with the difference between the glycoforms being the Hex elongating the *O*-fucose and not one of the *C*-Man. Red triangle: fucose; Green circle: mannose; Broken green circle: C-Man cross-ring fragment; White circle: Hex; Asterisk: cysteine carbamidomethylation

**Figure 2:**
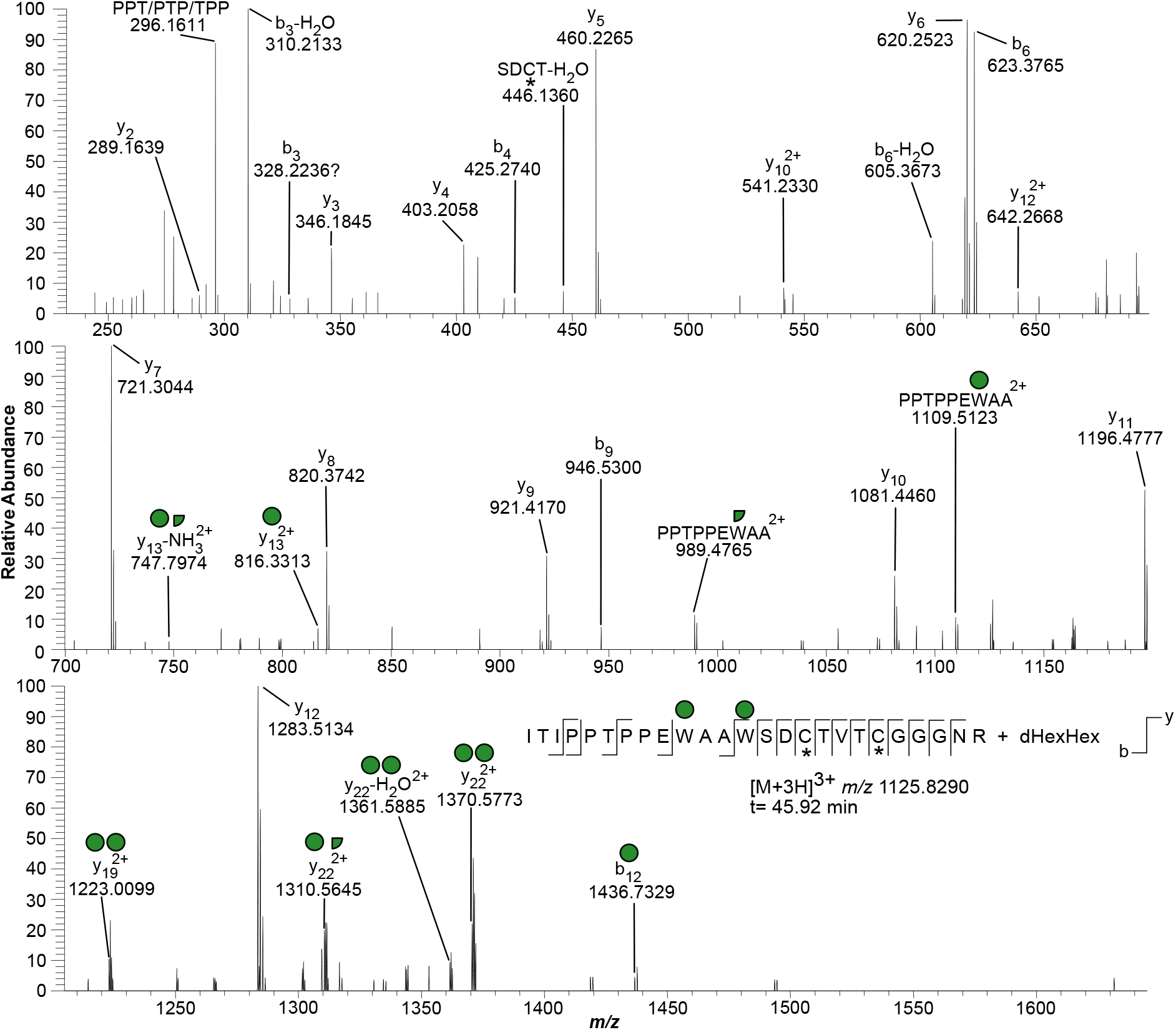
HCD MS/MS of an O-fucosylated and C-mannosylated glycopeptide from TSR3 of MIC2. HCD MS/MS with collision energy of 25 eV. Detection of the singly charged b11 and doubly charged y13, y13-NH_3_ ions allows positioning of C-Man residues to the first and second W of the WXXWXXC motif, respectively. Other ions of interest are the doubly charged y19 (two Hex residues) and y22 (two intact Hex(s) or one intact Hex plus one Hex -120.04 Da fragment). Red triangle: fucose; Green circle: mannose; White circle: Hex; Asterisk: cysteine carbamidomethylation.

**Figure 3:**
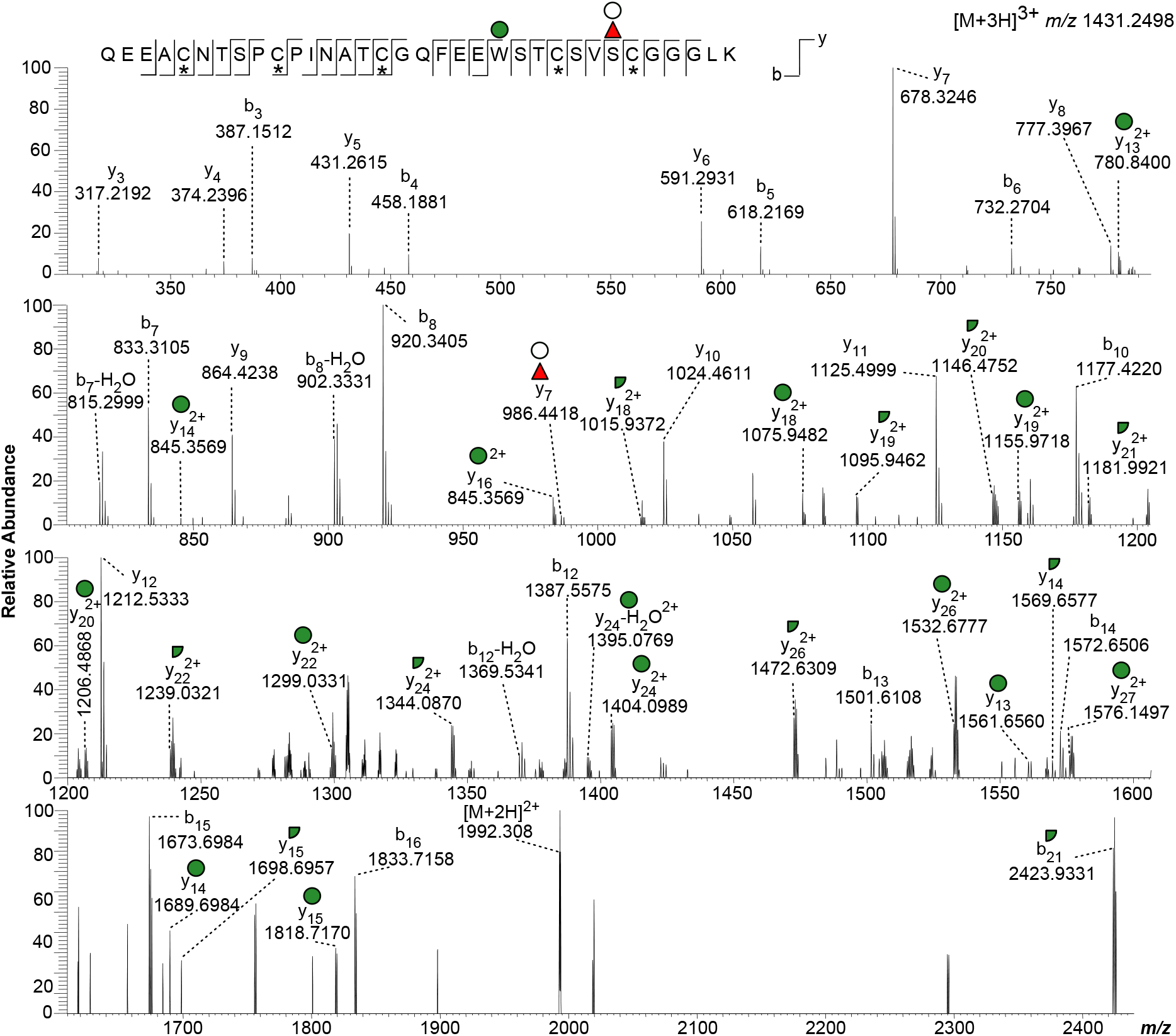
HCD MS/MS of an O-fucosylated and C-mannosylated glycopeptide from TSR4 of MIC2. HCD MS/MS spectrum using a collision energy of 20 eV. Detection of the Hex in the y-series ions only from y13 onwards, both on singly and doubly charged species, confirm positioning of the Hex on the Trp in the WXXC motif, consistent with C-mannosylation. The y7 + dHexHex ion support the presence of the disaccharide on Ser485, the predicted POFUT2 site. Red triangle: fucose; Green circle: mannose; White circle: Hex; Asterisk: cysteine carbamidomethylation.

A TSR1 semi-tryptic peptide [282]SVSCDGSQIR[294], modified with a dHex and a Hex, was observed as the [M+2H]^2+^ ion at *m/z* 737.3218 (Table 1). ETD fragmentation on an LTQ-Orbitrap resulted in three product ions that carried both Hex and dHex: z9, z10, and y10 (Fig. 1B). No other dHex and/or Hex containing product ions were observed. These observations allow assignment of Ser285, the predicted POFUT2 acceptor site, as the glycosylated residue and indicate that the two sugars compose a disaccharide modifying this serine (22). It should be noted that this analytical technique cannot define the hexose monosaccharide substituting the dHex.

For the MIC2 TSR3, we observed the semi-tryptic peptide [399]ITIPPTPPEWAAWSDCTVTCGGGNR[425] modified with a dHex and 3 Hex as a [M+3H]^3+^ species at *m/z* 1125.8290 (Table 1). HCD fragmentation using a low collision energy setting (10 eV) resulted in the detection of triply-charged product ions corresponding to loss of Hex from the precursor (*m/z* 1071.8176), loss of dHexHex (*m/z* 1023.1157), and subsequent loss of 120.04 Da, resulting from a cross-ring cleavage characteristic of C-mannose (*m/z* 983.1221) (Fig. 1C) (20). Increasing the collision energy to 15 eV, resulted in the detection of the ion species observed at 10 eV plus the precursor minus Hex, dHexHex, and two Hex fragments originating from cross-ring cleavages (*m/z* 843.0975). These results are consistent with peptide modification with two *C*-Man residues and a dHexHex disaccharide on the POFUT2 site. Additionally, loss of the Hex and then the dHex is consistent with the dHex as the monosaccharide that is attached to the protein. We also observed a lower abundance ion corresponding to loss of dHex from the precursor (*m/z* 1077.1379, Fig 1C). This ion could indicate either a minor glycoform where the Hex is the monosaccharide attached to the protein or a rearrangement of the glycan during the mass spectrometry analysis (34). Since rearrangement should be observed for other O-fucosylated TSRs, we analyzed tryptic peptides from human thrombospondin 1 (TSP1). HCD MS/MS of a semi-tryptic peptide from the TSR2 of TSP1, using a collision energy of 15 eV, resulted in a product ion ([M+2H]^2+^ *m/z* 663.8168) corresponding to the precursor after loss of a dHex (Fig. S2). This observation is consistent with the M-dHex ion resulting from rearrangement of the dHexHex. Confirmation of the presence of Hex residues on each of the two Trp in the WXXWXXC motif was obtained by HCD fragmentation using higher collision energies (20 and 25 eV) which resulted in almost complete elimination of the precursor ion (Fig. 2). In particular, detection of the doubly-charged y13 and y13-NH_3_ ions allowed positioning of one C-Man at the second W of the WXXWXXC motif; while the singly-charged b11 was diagnostic of the presence of another C-Man on the first W of the motif. These assignments were supported by the detection of the doubly charged ions y19 (two intact Hex residues, *m/z* 1223.0099) and y22 (two intact Hex(s), *m/z* 1370.5773), accompanied by product ions that arise via loss of water or 120.04 Da from y22 (*m/z* 1361.5885 and *m/z* 1310.5645, respectively) (Fig. 2).

Two peptides were observed for the MIC2 TSR4: [457]QEEACNTSPCPINATCGQFEEWSTCSVS CGGGLK [492] as a [M+3H]^3+^ species, modified with dHex and 2 Hex, at *m/z* 1431.2498 and a smaller semi-tryptic peptide [482]SVSCGGGLK[492] as a [M+2H]^2+^ species, modified with a dHexHex, at *m/z* 586.7708 (Table 1). ETD analysis of the smaller semi-tryptic peptide (*m/z* 586.7708) showed the presence of the z7 product ion carrying both Hex and dHex. This observation allowed assignment of Ser485, the predicted POFUT2 acceptor site, as the residue modified by dHexHex (Fig. 1D). Additionally, HCD fragmentation using low collision energy setting (10 eV) of the [M+3H]^3+^ species showed ions corresponding to the precursor after loss of an intact Hex (*m/z* 1377.2067), a Hex and dHex (*m/z* 1328.5223), or a dHexHex and a 120.04 Da from a Hex cross-ring cleavage (*m/z* 1288.5155) (Fig. S3). These results are consistent with modification of the Trp on the WXXC motif with one Hex and a dHexHex disaccharide modifying the *O*-fucosylation site. HCD fragmentation using higher collision energy (20 eV) allowed confirmation of Trp modification by a Hex upon detection of the sugar in the y-series ions only from y13 onwards, both as singly and doubly charged species (Fig. 3). HCD fragmentation also identified a y7 + dHexHex ion, consistent with the presence of the disaccharide on Ser485.

Lastly, multiple peptides containing the *C*-mannosylation and *O*-fucosylation sites were observed for MIC2 TSR5. The fully tryptic peptide [533]PGEWAEWGECSVTCGDGVR[553] modified by dHexHex2 was observed as a [M+2H]^2+^ at *m/z* 1311.5304 and the semi-tryptic peptide with one missed cleavage [526]ACPVDERPGEWAEWGECSVTCGDGVR[553] also carrying dHexHex_2_ as [M+3H]^3+^ at *m/z* 1150.4752 (Table 1). HCD MS/MS for the glycopeptide at *m/z* 1311.5304 has been previously shown in (25); it demonstrated that both Trp residues on the WXXWXXC motif are modified with Hex. The previously published HCD analysis also detected a y7 + dHex ion, consistent with the presence of deoxyhexose on Thr546 (25). In this study, we observed the fully tryptic peptide [533]PGEWAEWGECSVTCGDGVR[553] as [M+3H]^3+^ *m/z* 928.7054, consistent with its modification by dHex and three Hex residues. Additionally, we observed the tryptic peptide (one missed cleavage) [517]TETVTCNPQACPVDERPGEWAEWGECS VTCGDGVR[553] modified with dHexHex_2_ ([M+4H]^4+^ *m/z* 1120.7179) and with dHexHex_3_ ([M+4H]^4+^ *m/z* 1161.2313). HCD fragmentation using low collision energy (10 eV) was performed on both species. The identified fragment ions are consistent with the difference between these two glycoforms being a Hex on the *O*-fucosylation site since the m/z value measured for the most abundant ion, corresponding to the precursor minus the *O*-linked sugar, was the same within 10 ppm: *m/z* 1084.2029 and 1084.1923, respectively (Fig. 1E). Finally, a semi-tryptic peptide containing the TSR5 O-fucosylation site, [543]SVTCGDGVR[553], was also observed in two different glycoforms: modified with only a dHex ([M+2H]^2+^ *m/z* 548.7500) and with dHexHex ([M+2H]^2+^ *m/z* 629.7768).

In conclusion, all four MIC2 TSRs with POFUT2 acceptor sites are modified by a dHexHex disaccharide, consistent with FucGlc, and TSR5 can also be modified by only a dHex. Furthermore, Trp residues on TSR3 and 4, in addition to the already described TSR5 (25), are modified with *C*-mannose.

### Identification and gene disruption of *T. gondii* orthologues of host POFUT2 and GDP-fucose transporter

The *T. gondii* genome encodes for a putative POFUT2 (TGGT1_273550), which belongs to the CAZy GT68 family of glycosyltransferases that includes *Plasmodium* and human POFUT2 (Fig. S4) (35). Three conserved motifs, which are shared with the members of the α1,2-, α1,6- and O-FTs superfamily (*black boxes*), can be identified, including Motif I that forms part of the substrate binding site (36). A GT68-specific region (aa 229-357) includes the catalytic glutamic acid and some of the residues involved in TSR recognition (*gray boxes* in Fig. S4) (36, 37). *T. gondii* POFUT2 also has a hydrophobic N-terminal sequence (aa 1-180) which is absent from the other POFUT2s identified to date (Fig S4). Using CRISPR/Cas9 gene editing (38) in the RH ΔKu80 strain, which is deficient in the non-homologous end-joining DNA repair system (39), we added a sequence encoding a C-terminal 3xMYC tag to the endogenous *pofut2* locus (Fig. 4A and Table S2). Insertion of the 3xMYC tag at the correct locus was confirmed by PCR (Fig. 4B and Table S2) and the parasites were analyzed by immunofluorescence at 24 h post-electroporation to look for MYC-positive vacuoles. *T. gondii* POFUT2-3xMYC has a similar distribution to P30HDEL-YFP (40) (Fig. 4C), showing that POFUT2 is expressed in tachyzoites, and, for the most part, localizes to the endoplasmic reticulum, as expected (22).

**Figure 4:**
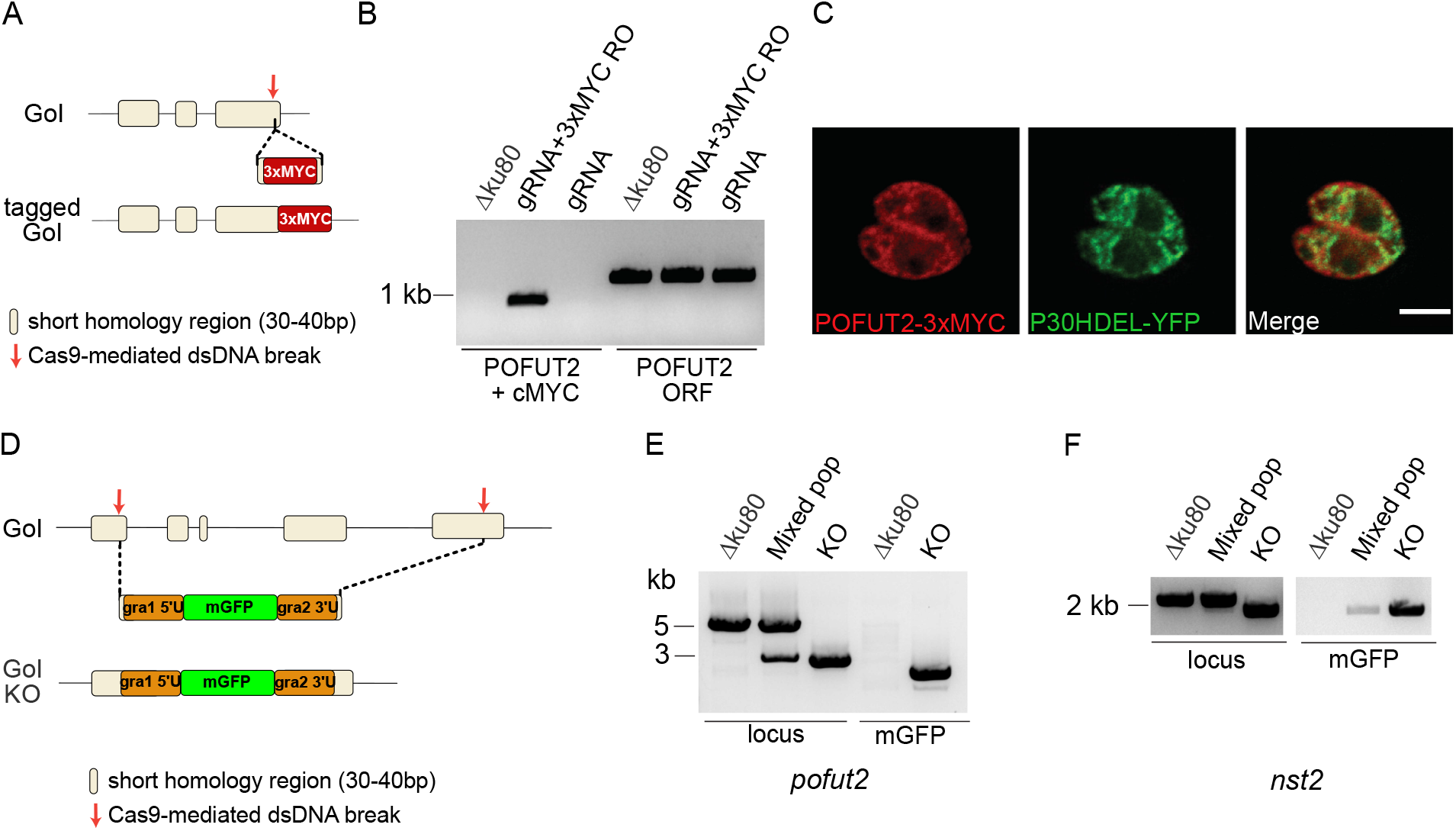
Localization of POFUT2 and generation of *Apofut2* and *\nst2* mutants. A. Schematic representation of the strategy used to *in situ* tag *pofut2*. Cas9 was directed to excise the *pofut2* gene 20 bp upstream of the stop codon and an oligonucleotide, flanked by about 50 bp of homology sequence to the break site, was provided as template for the homologous recombination. B. DNA was extracted at 72 h post electroporation and analyzed by PCR to show integration of the 3xMYC tag in the correct locus (ORF Fwd + cMYC rev) only when cells were electroplated with the CRISPR/Cas9 construct directing to the *pofut2* locus and the 3xMYC recombination sequence. A control reaction amplifying a fragment of the *pofut2* ORF was also performed (OFT Fwd + Rev). *C*. IFA of RH AKu80 tachyzoites expressing *pofut2* tagged at the C-terminus with a 3xMYC tag and an ER marker (p30HDEL-YFP) shows partial colocalization. Scale bar: 2 μm. *D*. Schematic representation of the strategy used to generate the *pofut2* and *nst2* gene disruptions. Cas9 was directed to excise each gene in the first and last exon. An mGFP-expressing cassette, flanked by about 30 bp of homology sequence to each break site, was provided as template for the homologous recombination. *E-F*. Genomic DNA was extracted after a first enrichment for mGFP-positive cells (mixed pop) and after the final cloning step (KO). PCR analyses show substitution of the wild type ORF with the mGFP cassette in both strains. Mixed pop: mixed population.

Transport of the activated sugar GDP-Fuc from the cytosol to the lumen of the ER is required for POFUT2 activity. A BLASTP search, using the human Golgi GDP-Fuc transporter as template, identified a single putative GDP-sugar transporter, TGGT1_267730 or NST2. There are 10 predicted transmembrane helices (41) and conserved nucleotide and sugar binding motifs (*black dashed boxes*) at the C-terminus (42) (Fig. S5).

Using CRISPR/Cas9 in the RH AKu80 strain, the coding regions of *T. gondii pofut2* and *nst2* were disrupted, as described in Fig. 4D. Briefly, a cassette expressing mGFP, under the *gra1* promoter (43), was inserted in either locus (Fig. 4D and Table S2). After selection for mGFP-positive cells, clones were obtained by limiting dilution. Integration at the correct locus was verified by diagnostic PCR for both *Δpofut2* (Fig. 4E) and *Δnst2* (Fig. 4F) clones (see Table S2 for primer sequences).

### MIC2 is not *O*-fucosylated in *Apofut2* and *Anst2*

To determine the glycosylation status of the mutants, we used an antibody against the disaccharide FucGlc. To generate the antibody, chickens were immunized with Glc-ß-1,3-Fuc-a-KLH (16) as antigen (Fig. 5A and Supporting Information), and the disaccharide-specific polyclonal IgY were purified from the antibody fraction via Glc-ß-1,3-Fuc-a-agarose beads. To analyze the anti-GlcFuc IgY specificity, the disaccharide and the two monosaccharides (Fuc and Glc) were conjugated to BSA. The resulting neoglycoproteins (BSA-GlcFuc, BSA-Fuc and BSA-Glc) were then tested by ELISA and Western blot. Antibody titration by ELISA identified 1 μg/ml as the concentration with the best signal to noise ratio (Fig. 5B and S6). Western blot analysis also confirmed the antibody specificity for BSA-GlcFuc, although a minor reactivity to BSA-Fuc was also observed (Fig. 5C).

**Figure 5:**
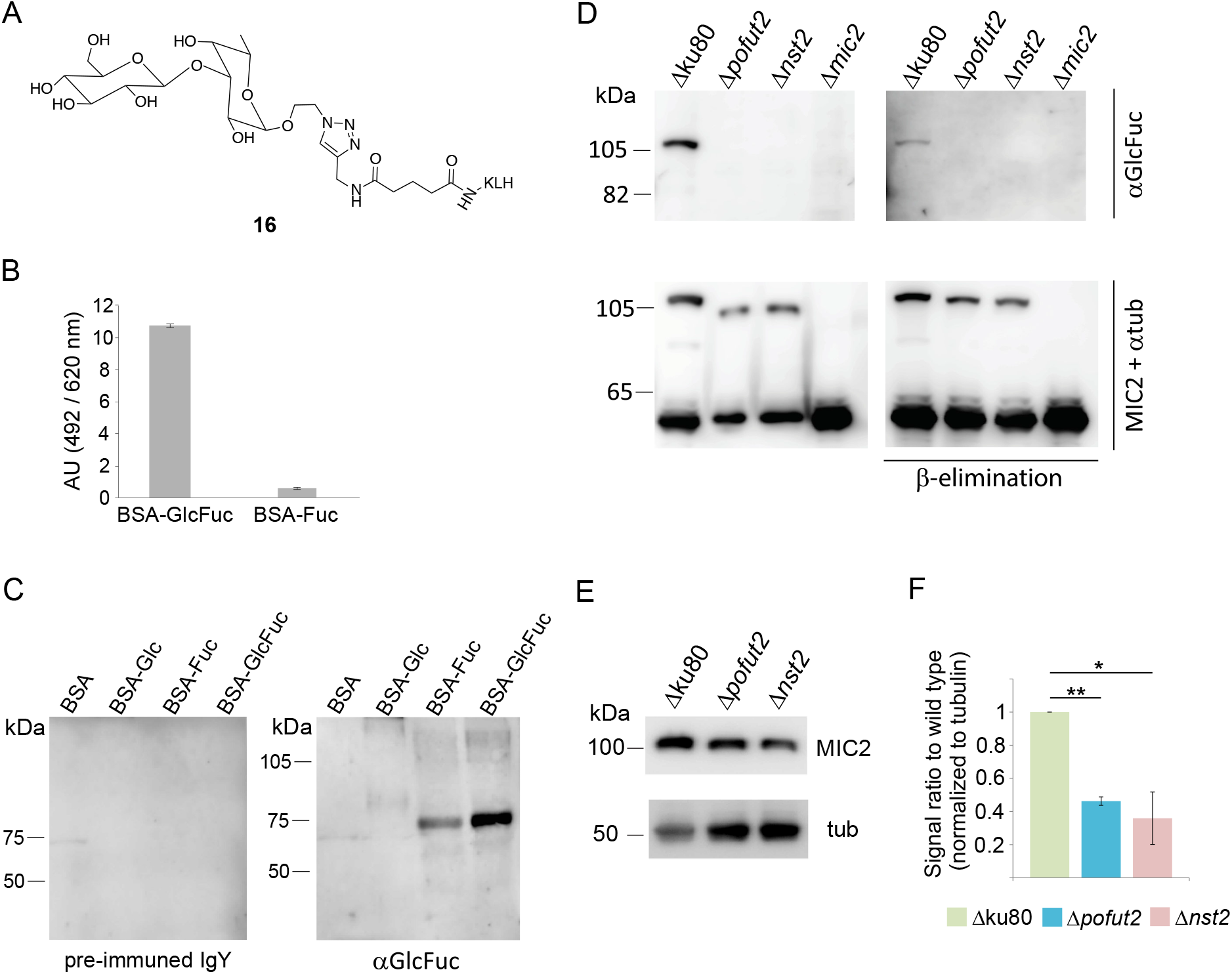
MIC2 is not O-fucosylated in either *Apofut2* and *\nst2* strains. A. Structure of Glc-ß-1,3- Fuc-a-KLH antigen B. ELISA assay of the purified anti-GlcFuc antibody (1 μg/ml) shows the specificity of the polyclonal IgY for BSA-GlcFuc versus BSA-Fuc. C. Western-blot testing of purified anti-GlcFuc antibody confirms its specificity for BSA modified with the GlcFuc disaccharide. No reactivity is observed probing with the pre-immune IgY fraction. *D*. Western blot analysis with anti-FucGlc antibody identifies a ß-elimination sensitive band in wild type corresponding to MIC2. No reactivity is shown in all three knockouts analyzed. MIC2 and tubulin controls are also shown. *E*. Western blot of tachyzoites cell lysate with monoclonal anti-MIC2 shows reduced levels of cellular MIC2 in both knockouts. *F*. MIC2 levels were normalized to tubulin and the average of three biological repeats is shown. Student t test was used to compare the samples and significant differences are marked with * (0.05<p<0.01) and ** (p=0.001).

When total cell lysates from the parental strain and the *O*-fucosylation mutants were analyzed by Western blot using the antibody against GlcFuc, a ß-elimination-sensitive band at an apparent MW of about 110 kDa was identified in wild type lysate, but not in the *Δpofut2* and *Δnst2* lysates (Fig. 5D). Additionally, the anti-GlcFuc antibody did not bind to *Δmic2* lysate, confirming that the band observed in wild type is indeed MIC2 and that MIC2 TSRs are not being O-fucosylated in the two O-fucosylation mutants (Fig. 5D). These results also indicate that the dHexHex identified by mass spectrometry is indeed GlcFuc.

Using the fucose-specific *Aleuria aurantia* lectin (AAL), we have previously described *O*-fucosylated proteins at the nuclear periphery of *T. gondii* (44). To assess the role of TgPOFUT2 on nuclear *O*-fucosylation, we analyzed the mutant parasites by AAL immunofluorescence. No effect on AAL binding was observed upon knockout of *pofut2*, consistent with nuclear O-fucosylation being mediated by a different glycosyltransferase (Fig. S7).

Loss of *O*-fucosylation resulted in a decrease of more than 50% in cellular MIC2 levels in both mutants, as determined by Western blot analysis (Fig. 5E and 5F). This result suggests that *O*-fucosylation may be important for protein folding or processing of MIC2, since non-fucosylated MIC2 is not being processed as efficiently as wild type MIC2.

### Absence of *O*-fucosylation results in defective MIC2 localization to the micronemes

Localization of MIC2 was analyzed by immunofluorescence microscopy. As shown in the left panel of Fig. 6A, 12-15% of vacuoles in both *Δpofut2* and *Δnst2* show an accumulation of MIC2 to the early/mid secretory pathway (about 15% of vacuoles for *Δpofut2* and 12% for *Δnst2*, Fig. 6B). The same abnormal accumulation was observed for M2AP (Fig 6A middle panel), but not for AMA4 (Fig. 6A, right panel), a protein with no TSRs and no reported strong association with MIC2. These observations suggest we are observing an effect specific to O-fucosylated proteins and not to all micronemal proteins. In both *Δpofut2* and *Δnst2* strain, transiently expressed GRASP55-mRFP, a *cis* Golgi marker (40), partially co-localizes with the fraction of MIC2 that accumulates in the early/mid secretory pathway (Fig. 6C). IFA also suggests that GRASP55 distribution is aberrant in these vacuoles, as it looks more diffused and not like the observed cisternae-shaped structure observed in the parental strain (Fig. 6C). Both the reduction of MIC2 levels and its defective localization are consistent with a role for *O*-fucosylation in protein folding and secretion, as has been described in humans and *P. falciparum* (23, 32).

**Figure 6:**
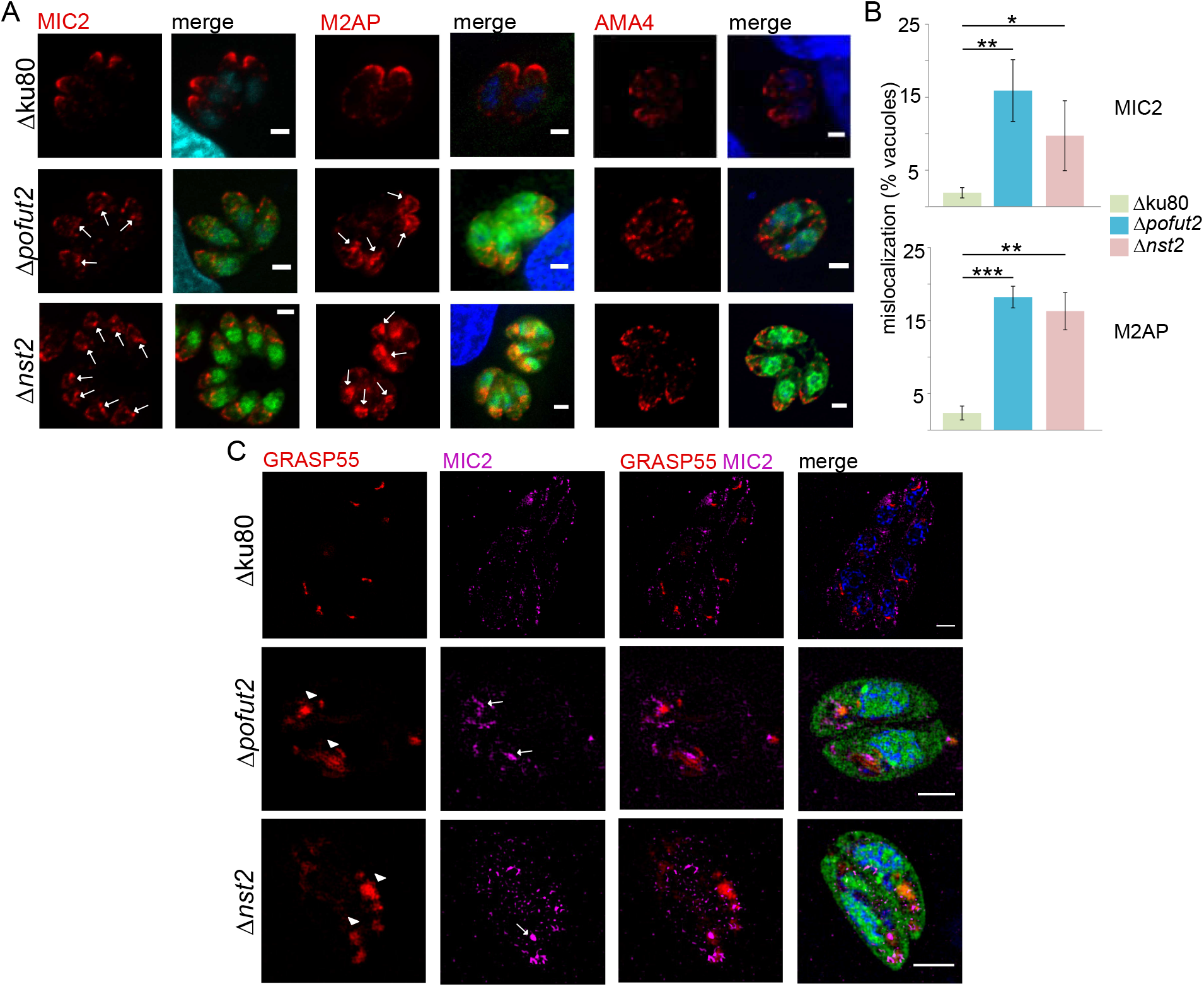
MIC2 is mislocalized in both *Apofut2* or *Anst2* strains. A. MIC2, and consequently M2AP, localization is affected in the KO with both proteins accumulating in the early secretory pathway in a fraction of the vacuoles (*white arrows*). This is not a general microneme defect as AMA4 location is unaffected. Merge: antibody staining (red), DAPI (blue), and mGFP (green) for the O-fucosylation mutants. B. Quantification of the percentage of vacuoles with abnormal MIC2 and M2AP localization. The average of three biological repeats ± SD is shown. Student t test was used to compare the samples and significant differences are marked with * (0.05<p<0.01), ** (0.01<p<0.0001), and *** (p<0.0001).C. Parasites were transiently electroporated with GRASP55-mRFP and analyzed by structured illumination microscopy. Mislocalized MIC2 partially co-localizes with this *cis* Golgi-marker and that the GRASP55 localization itself is aberrant (*arrow heads). White arrows* mark accumulation of MIC2 in the early/mid secretory pathway. Merge: MIC2, GRASP55-RFP, mGFP, and DAPI. Scale bars: 2 μm.

### Phenotypic analyses show a defect in attachment and invasion

The genome-wide CRISPR/Cas9 screen that has been performed in *T. gondii* (45) identified both *pofut2* and *nst2* as non-essential genes (phenotypic scores of -0.34 and +1.23, respectively). Here we found that both *Δpofut2* and *Δnst2* displayed a 40% reduction in the number of plaques compared to the parental strain (Fig. 7A and 7B), while no difference between mutants and wild type was observed in the average area of the plaques (Fig. S8). These observations are usually indicative of a defect in the parasite’s ability to either invade or egress from the host cells.

**Figure 7:**
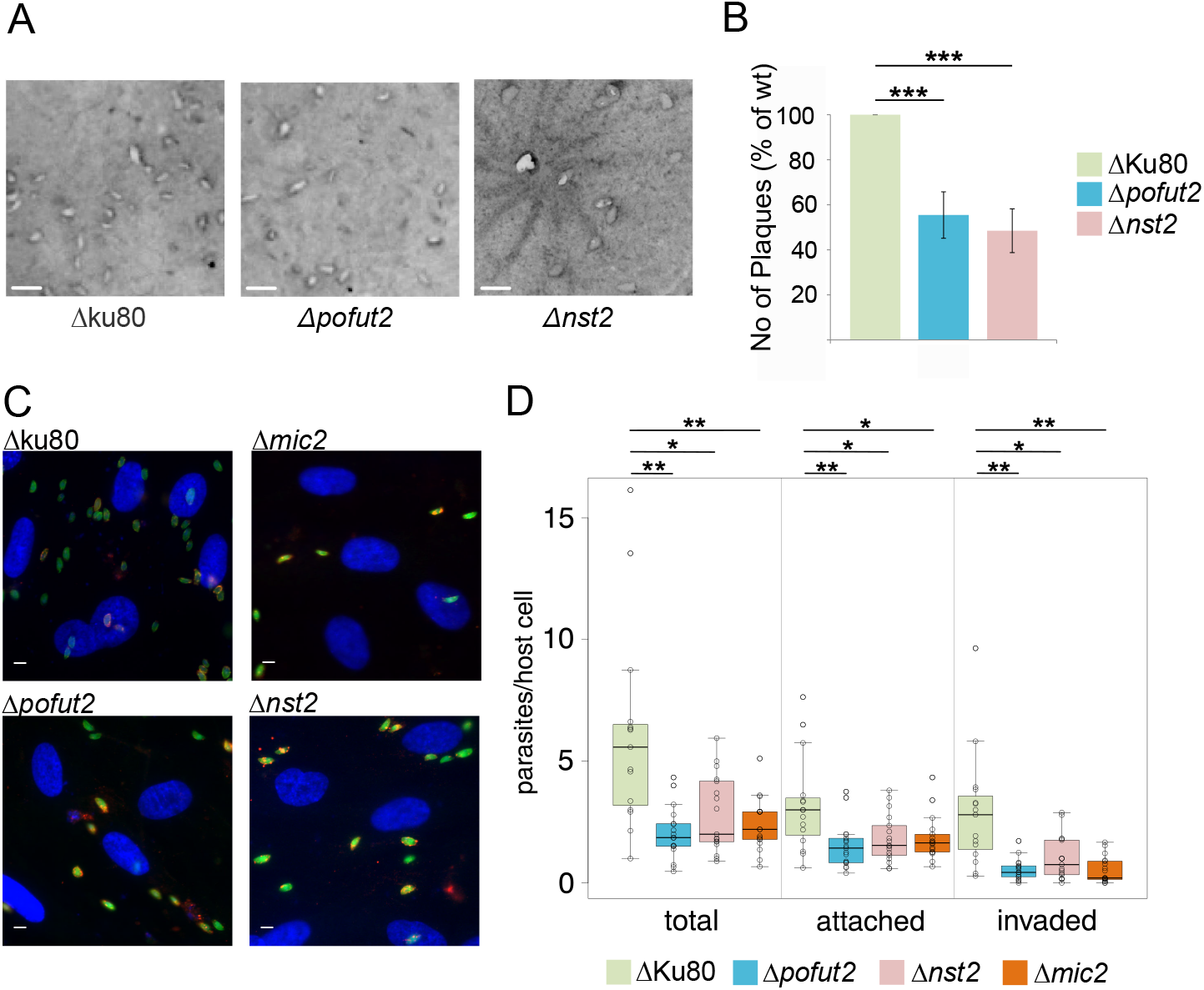
Growth, attachment, and invasion are affected in *Apofut2* and Anst2. A. Representative images from the plaque assay. Scale bars: 2 mm. B. Quantification of the number of plaques observed shows a 40% reduction is growth in the KOs compared to the parental strain. The average of four biological repeats ± SD is shown. *** (p<0.0001) C. Representative images from the red/green invasion assay. Extracellular parasites were labeled with anti-SAG1 and are shown in red. After permeabilization, all parasites were labeled with anti-ß-tubulin (in green). As a result, attached parasites look yellow (red + green) and invaded parasites are green. B. Boxplot for the red/green assay, showing the number of parasites counted in each field and normalized to the number of host cells nuclei in the same field. For each cell line, the number of total parasites, and attached (red + green) or invaded (green) parasites are shown. Student t test was used to compare the samples and significant differences are marked with * (0.05<p<0.01) or ** (p<0.01). All *p* values are reported in Table S3. All quantifications are given as the average of at least three biological repeats ± SD. Scale bars: 5 μm

Phenotypic assays were performed to analyze the knockouts’ ability to attach, invade and egress. *Δpofut2* and *Δnst2* were compared with the parental strain and the recently published *Δmic2* (13), as the data presented here indicate MIC2 is the most abundant O-fucosylated protein in tachyzoites and is incorrectly processed in both knockouts. Attachment and invasion were assessed via the previously described red/green assay (Fig. 7C) (46). A significant reduction is the total number of parasites that attached to or invaded the host cells was observed in all three knockout cell lines, compared to the RH Aku80 strain (Fig. 7D). Additionally, no statistical difference was observed between *Δpofut2* or *Δnst2* and *Δmic2*, suggesting lack of O-fucosylation is sufficient to reproduce the attachment/invasion defect observed when MIC2 is not being synthesized (Table S3). Furthermore, there was no statistical difference between *pofut2* and *nst2* knockouts.

Egress was stimulated with addition of the calcium ionophore A23178, as described in the Experimental Procedures, and intact, permeabilized or completely egressed parasitophorous vacuoles (PVs) were quantified (Fig. 8A). As shown in Fig. 8B and Table 2, all three knockouts had a significantly larger proportion of PVs that were just permeabilized, compared to the RH *Δ*ku80 strain. However, the egress defect in *Δmic2* was found to be significantly more severe than in either *Δpofut2* or *Δnst2* (Table 2). The increased egress defect in the *Δmic2* strain may explain the markedly reduced infectivity in culture of the *Δmic2* strain, reported in (13), versus the more modest reduction in the *Δpofut2* and *Δnst2* strains shown here.

**Table 2:** Comparison of egress phenotypes between wild type and the knockout cell lines analyzed in this study.

| Phenotype | $\Delta ku80$ vs.<br>$\Delta pofut2$ | $\Delta ku80$ vs.<br>$\Delta nst2$ | $\Delta ku80$ vs.<br>$\Delta mic2$ | $\Delta pofut2$ vs.<br>$\Delta nst2$ | $\Delta pofut2$ vs.<br>$\Delta mic2$ | $\Delta nst2$ vs.<br>$\Delta mic2$ |
| --- | --- | --- | --- | --- | --- | --- |
| Egressed | 0.04 | 0.0007 | $8.1 \times 10^{-15}$ | n.s. (0.18) | $2.3 \times 10^{-10}$ | $1.8 \times 10^{-8}$ |
| Permeabilized | 0.007 | 0.0001 | $1.8 \times 10^{-15}$ | n.s. (0.13) | $9.5 \times 10^{-11}$ | $8.8 \times 10^{-8}$ |
| Intact PSV | n.s. (0.92) | n.s. (0.88) | n.s. (0.62) | n.s. (0.95) | n.s. (0.56) | n.s. (0.53) |

n.s.: not significant

**Figure 8:**
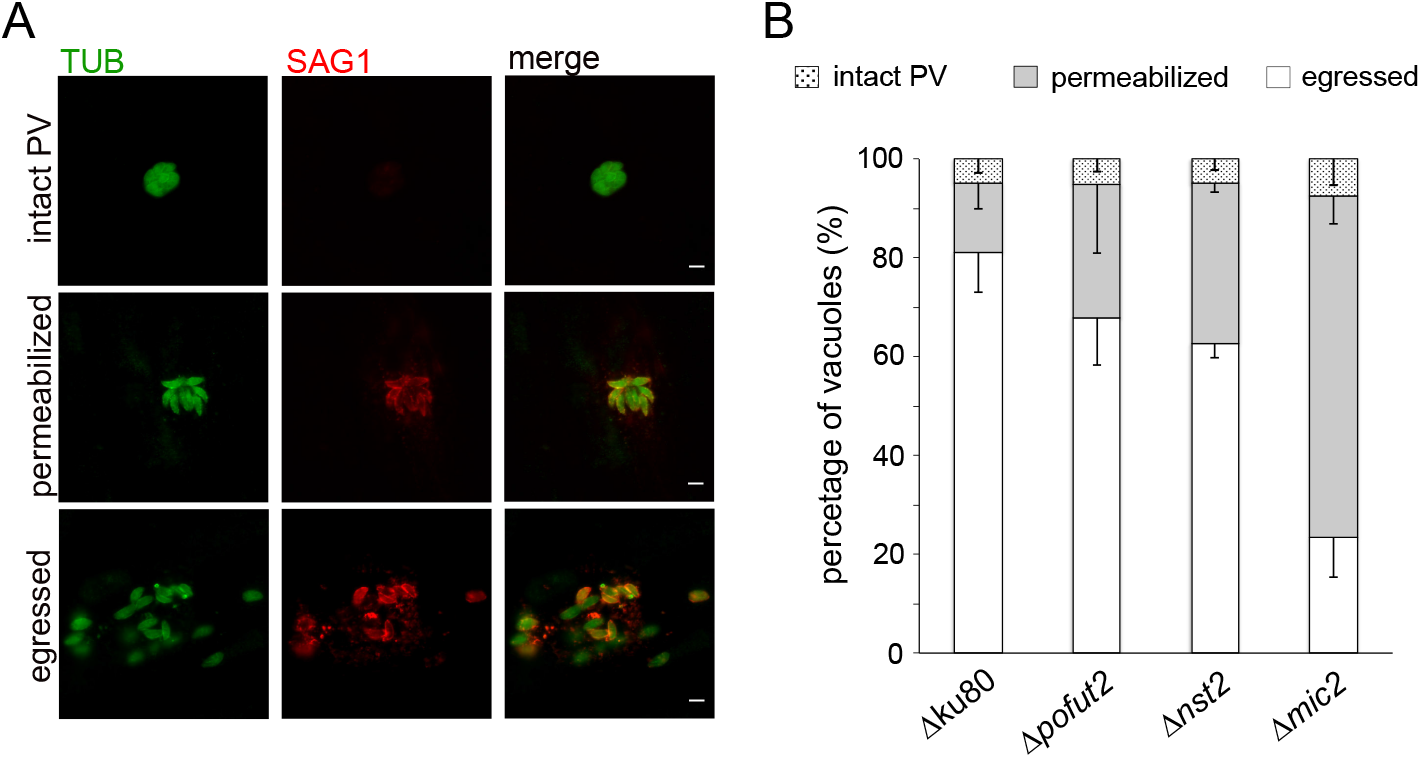
Egress defect in *Apofut2* and *Anst2* is not as severe as in the *Amic2*. A. Representative images showing the different phenotypes observed in the assay. Extracellular parasites were labeled with anti-SAG1 (in red). After permeabilization, all parasites were labeled with anti-ß-tubulin (in green). As a result, parasites in intact vacuoles are green, while parasites in permeabilized vacuoles or that have egressed are labeled in green and red. B. Quantification of the egress phenotypes. Quantifications are given as the average of at least three biological repeats ± SD. Student t test was used to compare the samples and all*p* values are reported in Table 2. Scale bars: 5 μm.

## Discussion

MIC2 is a well-characterized *T. gondii* virulence factor. This study adds to our understanding of MIC2 structure and function by characterizing the glycosylation on its TSRs and describing the importance of *O*-fucosylation on MIC2 processing and, consequently, the parasites ability to invade.

Mass spectrometry analysis, using both HCD and ETD, showed that, in MIC2, all four TSRs containing the O-fucosylation motif (TSR1, 3,4, and 5) are modified by *O*-FucGlc. Two other TSRs (TSR3 and 4), in addition to TSR5 (25), are also modified with *C*-Man. TSR5 was observed *in vivo* in two glycoforms indicating heterogeneity at the *O*-fucosylation site, which is modified with either Fuc or FucGlc. While ETD-based fragmentation is a very effective technique to characterize glycan modifications and identify the glycosylated amino acids, it is not able to distinguish between isobaric sugars, *e.g*. glucose vs. galactose. The evidence which indicates the disaccharide is FucGlc, is the binding by Western blot of an antibody generated here against the Glcß1,3Fuc disaccharide. Consistent with this observation, a putative B3GLCT, the enzyme that adds Glc to elongate the Fuc on TSRs, is also present in the *T. gondii* genome (TGGT1_239752).

MIC2 was the only one of four predicted POFUT2 substrates that was identified in our mass spectrometry analysis of secreted proteins, as also reported in (25). While it is possible that MIC2 is the only tachyzoite protein with TSRs modified by *O*-Fuc and C-Man, it is more likely that the failure to detect other targets was due to either their low abundance or lack of secretion. TGGT1_223480, for example, is predicted to have higher expression in unsporulated oocysts (47) and could be the protein responsible for the previously observed staining of oocyst walls by the fucose-specific AAL (44), providing *O*-Fuc is not capped in this life stage.

Eukaryotic POFUT2s have been described as soluble ER lumen proteins (22), and immunofluorescence after *in situ* tagging of *T. gondii* POFUT2 indicates the ER localization is conserved in the parasite. However *in silico* analyses suggested a different topology, as four putative transmembrane domains (TM) are predicted in the hydrophobic N-terminal region of *T. gondii* POFUT2. Attempts at recombinantly express POFUT2 either plus or minus the N-terminal domain or with only the fourth TM in CHO cells, as previously done for DPY19 (25), did not result in active protein (data not shown). Even in the absence of *in vitro* activity data, loss of O-fucosylation on MIC2 in the *Δpofut2* strongly supports the model that this enzyme is the *O*-fucosyltransferase that modifies TSRs in the parasite.

*T. gondii* is predicted to synthesize both GDP-Man and GDP-Fuc, but only one putative GDP-sugar transporter, NST2, can be identified in its genome. Consistent with this observation, the C-mannosyltransferase DPY-19 (25) utilizes Dol-P-Man as a sugar donor (26), and no putative Golgi mannosyl-or fucosyl-transferases can be identified. Additionally, no putative POFUT1, the enzyme responsible for *O*-fucosylation of endothelial growth factor-like domains (48), is present in the parasite genome, even though *T. gondii* is predicted to encode for multiple EGF domain-containing proteins (49). The lack of *O*-fucosylation in *Δnst2* parasites and the similar phenotype between *Δpofut2* and *Δnst2* mutants support the identification of NST2 as *T. gondii* GDP-Fuc transporter and suggests that in the parasite secretory system GDP-Fuc is primarily used for *O*-fucosylation by POFUT2. Identification of the GDP-Fuc transporter involved in TSRs and EGFs O-fucosylation has proven more difficult in metazoans with various studies suggesting that the players involved in GDP-Fuc transport into the ER are not as well conserved as the O-fucosyltransferases themselves (50-52).

Studies on mammalian POFUT2 indicate that this enzyme recognizes already folded TSRs and defects in its activity affect the folding dynamics or the stabilization of TSR-containing proteins (37, 53). Our observations in *T. gondii* are consistent with this model. Although we detect non-fucosylated MIC2 that is correctly trafficked (and presumably folded), we also observe in a subset of vacuoles a clear accumulation of likely misfolded MIC2 in the early/mid secretory pathway. Aberrant GRASP55 staining, which is also observed in this subpopulation of vacuoles, suggests that accumulation of misfolded MIC2 causes a disruption in the Golgi structure. Our results also suggest that the M2AP/MIC2 complex is not affected in either *Δpofut2* or *Δnst2*, as we observed mislocalization of MIC2 and M2AP to the early/mid secretory pathway in a statistically comparable number of vacuoles. This is unlike the *Δmic2*, where M2AP is redirected to the constitutive secretion pathway (13). Previous work has shown that M2AP binds MIC2 TSR6 through its galectin-like domain, and structural studies indicated the recognition is due to protein-protein, not carbohydrate-protein, interactions (16). While MIC2 TSR6 contains a C-mannosylation motif, we observed it exclusively in its unglycosylated form, supporting the protein-protein interaction model.

Comparison between the two *O*-fucosylation-deficient strains generated in this study and the previously described *Δmic2* (13) indicates that glycosylation of TSRs has the strongest effects on attachment and invasion (Fig. 9). This result suggests a role for MIC2 TSRs in the attachment and/or invasion processes, consistent with the known role of these domains in cell-cell interaction (17). These results are in agreement with the recently described *pofut2* knockout in *Plasmodium falciparum*, which also resulted in reduced invasion efficiency by sporozoites (32).

**Figure 9:**
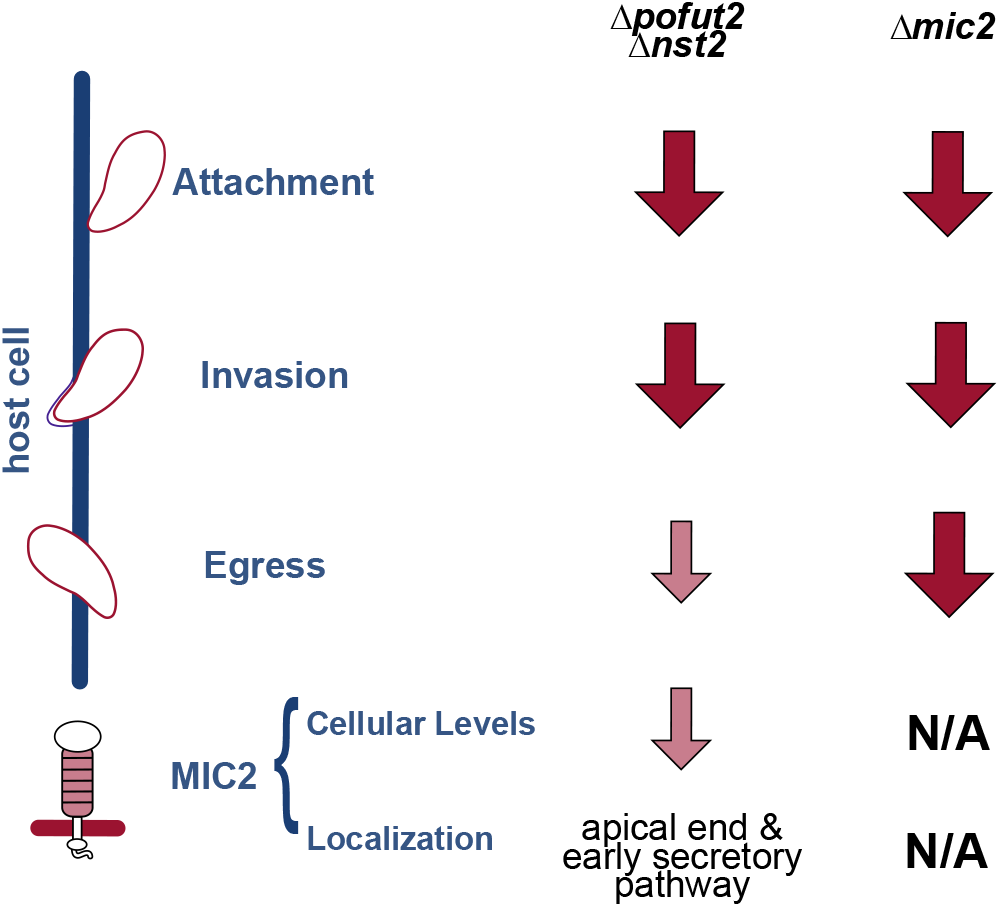
Comparison of *mic2* and O-fucosylation KOs. Schematic representation of differences and similarities between *Δmic2* and *Δpofut2* and *Δnst2*. Disruption of O-fucosylation is sufficient to reproduce the attachment and invasion defects observed when MIC2 is not being synthesized. Conversely, the parasite ability to egress is much more compromised in the *Δmic2* versus *Δpofut2* and *Δnst2*. N/A: not applicable.

This work complements the recent studies that have looked at the role of fucosylation in *T. gondii* biology. Fucose has been previously described in this parasite as part of two uncommon nucleocytosolic glycosylation pathways: modification of the E3-ubiquitin ligase adaptor Skp1 by a fucose-containing pentasaccharide (54, 55), and an *O*-fucosylation pathway that modifies Ser/Thr on proteins that accumulate at the nuclear periphery (44). In this study we add a role for fucose in folding or stabilization of TSRs in *T. gondii*, a function conserved from *Plasmodium* to metazoans (23, 31, 32).

## Experimental Procedures

### In silic oanalyses

Human POFUT2 (UniProt # Q9Y2G5-3) and GDP-Fucose transporter (UniProt # Q96A29-1) protein sequences were used as templates to mine the *T. gondii* genome (ToxoDB.org, (47). The hits from these searches were confirmed by using the hits themselves as templates for searches on the NCBI BLASTP engine. Alignments were performed using ClustalOmega (56) and were colored by percentage identity and edited using Jalview (57). To identify the putative POFUT2 protein acceptors, a protein motif search was performed on ToxoDB for C^1^X2-3(S/T)C^2^X2G (47). The resulting hits were manually verified to confirm the proteins had a predicted signal peptide and the motif was indeed part of a TSR. The identified proteins are listed in Table S1.

### *Toxoplasma gondii* culture and manipulation

Cell culture of *T. gondii* tachyzoites was performed as previously described (39, 58). The *Δmic2* cell line was a kind gift of Dr. Markus Meissner, Ludwig-Maximilians-University of Munich (13). RH Δku80 was used as parental strain for all *T. gondii* transgenic cell lines generated in this study (39). To generate the transgenic cell lines, the required amount of DNA (see below) was electroporated by adapting the protocol described in (45). Briefly, tachyzoites were washed in HHE buffer and resuspended in Cytomix to be 1.6x10^7^/ ml. Purified DNA that has been previously ethanol precipitated, was re-dissolved in Cytomix (supplemented with GSH, CaCl2 and ATP) and combined with 10^6^ tachyzoites in a 2 mm gap BTX cuvette (BTX Harvard Apparatus). Electroporation was performed using program X-014 on an Amaxa^®^ Nucleofactor^®^ II electroporator (Lonza).

### CRISPR/Cas9-mediated gene disruption

The approach to generate the *pofut2* (TGGT1_273550) and *nst2* (TGGT1_267730) knockouts is described in Figure 4D. The protospacers to direct Cas9 were cloned using oligonucleotides P24-25 (pofut2_Nterm), P26-27 (pofut2_Cterm), P28-29 (nst2_Nterm), and P30-31 (nst2_Cterm) (Table S2). Oligos were annealed and phosphorylated, and the resulting dsDNA was cloned in the BsaI digested pU6Universal plasmid (38) to generate the following plasmids: pU6_P2N, pU6_P2C, pU6_N2N, and pU6_N2C. To replace the gene of interest, the mGFP expressing cassette from pDHFR-Luc-mGFP(43) (gra1 5’UTRs-mGFP-gra2 3’UTRs) was amplified by PCR using primers containing about 30-40 bp homology sequence to the double stranded break sites (P20-21 and P22-23, TableS2) and cloned into pCR2.1TOPO (Invitrogen), generating pP2KO_mGFP and pN2KO_mGFP. The cassettes also contained blunt restriction sites (AfeI/BsrBI for *pofut2* and StuI/SnaBI for *nst2*) to allow their isolation from plasmid DNA, without adding or removing bases to the homology sequences. DNA was purified from pP2KO_mGFP and pN2KO_mGFP and the cassettes isolated by restriction digestion and gel purification. About 20 μg each of pU6gRNAs and at least 5x molar excess of purified recombination cassette were used for each electroporation. Cells were recovered in 15-cm culture dishes (Corning) and mGFP-positive plaques were isolated and transferred to 96-well plates for cloning by limiting dilution. Two to three rounds of cloning were required to isolate clonal populations of mGFP-positive parasites. DNA was extracted before and after cloning using DNAzol (Invitrogen) according to manufacturer’s instructions. Insertion of the cassette in the correct locus was verified by PCR using primers P34, P36, and P39 (*pofut2*) or P32, P33, and P34 (*nst2)* (Table S2).

### *In situ* tagging of POFUT2

The approach is shown in Fig. 4A. Briefly, pU6_P2C was used to generate a Cas9-mediated double stranded break near the stop codon of the *pofut2* coding sequence. A 258-bp recombination sequence (O1, TableS2) was designed to include a 3xMYC tag, flanked at both sides by about 50 bp of homology sequence and blunt restriction enzymes (MscI/PvuII) to allow release from the plasmid (pUC57-P2tRO), and sent for synthesis (GenScript). Additionally, a synonymous point mutation was introduced in O1 to remove the Cas9 PAM site after recombination (mutated base in *lower case* in Table S2) (38). About 20 μg of pU6_P2C and 25x molar excess of the O1 sequence were electroporated as described above in the presence or absence of 10 μg of ptubP30HDEL/sagCAT as an endoplasmic reticulum marker (40). Parasites were then allowed to infect human foreskin fibroblasts (HFF) monolayers on coverslips in 24-well plates for up to 72 h. Cells were fixed for immunofluorescence analysis at 24 h. post electroporation (p.e.). DNA was extracted 72 h p.e. and used to verify insertion of the 3xMYC tag at the correct locus by PCR (primers P36, P37, P38, and P40).

### Immunofluorescence analysis

Unless otherwise specified, intracellular tachyzoites were fixed in 4% paraformaldehyde (PFA) in phosphate buffer (PB) for 20 min at room temperature (RT). Permeabilization was performed in 0.25% TX-100 in phosphate-buffered saline (PBS) for 15 min at RT and it was followed by blocking in 3% bovine serum albumin (BSA) in PBS for 1 h, at RT. pGRASP55-mRPF/sagCAT(40) was a kind gift of Marc-Jan Gubbels, Boston College. Electroporation was performed as described above, using 20 μg of plasmid, and cells were fixed 24 h post-electroporation. Primary antibodies were used at the following concentrations: MAb anti-MIC2 6D10 1:500, rabbit anti-MIC2 1:500, rabbit anti-AMA4 1:1000, mouse anti-c-myc 9E10 (DHSB) 4 μg/ml and, rabbit anti-M2AP 1:500. Antibodies to MIC2, M2AP were kind gifts of Dr. Vern Carruthers, University of Michigan, while the antibody against AMA4 was a kind gift of Maryse Lebrun, UMR 5235 CNRS, Université de Montpellier. Goat AlexaFluor-conjugated anti-mouse and anti-rabbit antibodies (Molecular Probes) were used at a 1:800 dilution. *Aleuria aurantia* lectin (AAL) was purchased from Vector Labs and conjugated to AlexaFluor594 succinimidyl esters (Molecular Probes) following the manufacturer’s instructions. The final conjugate (AAL-AlexaFluor594) was used at 1:250. Cells were labeled with 1 μg/ml 4’,6-diamidino-2-phenylindole (DAPI) for 10 min at RT. Coverslips were mounted on glass slides using Vectashield (Vector Labs) as mounting medium and examined by deconvolution microscopy on a Zeiss AXIO inverted microscope with a Colibri LED. Images were collected at 0.2-μm optical sections with a Hamamutsu Orca-R2 camera and deconvolved using ZEN software (Zeiss). Images were further processed and single optical sections selected with Fiji (59). When counting the number of vacuoles with mislocalized MIC2 or M2AP, cells were observed at a total magnification of a 1000x and 100 vacuoles/biological repeat were counted. The average ± SD of three independent repeats is shown. A two-tailed, homoscedastic Student t test was used to calculate the p values. Super-resolution microscopy was performed on a Zeiss ELYRA microscope. Images were acquired with a 63x/1.4 oil immersion objective, 0.089 μm z sections at RT and processed for structured illumination (SIM) using ZEN software. Single optical sections were selected using Fiji (59).

### Affinity purification of polyclonal chicken antibody

Glc-ß-1,3-Fuc-a-KLH (16) (see Supporting Information) was send to Genway Biotech Inc. to generate chicken antibody using standard protocols. Crude chicken IgY (50 ml at 10 mg/ml) were centrifuged at 5000 x *g* for 5 min. The debris was removed and the clear IgY was incubated with agitation overnight with the disaccharide agarose beads (see Supporting Information). The mixture was loaded on a column and the flow through fraction collected. The column was washed with PBS buffer and eluted with 0.1 M glycine buffer pH 2. The flow through fraction was re-incubated with the agarose beads and elutions collected twice more to ensure maximum yield of the GlcFuc-antibody. The elutions were immediately neutralized with 1 M TrisHCl pH 9.0, pooled and concentrated to 1 mg/ml using Amicon ultra centrifugal filters (EMD Millipore) 3000 MWCO (60).

### Generation of BSA neoglycoproteins

The disaccharide (GlcFuc) or the monosaccharides (Glc and Fuc) with an azide linker group was reacted with the NHS-linker (12) (see Supporting Information) and BSA according to generate the BSA neoglycoproteins, using the same method for generating KLH-GlcFuc (see Supporting Information, synthesis of compound 16). The products were purified by dialysis in distilled water for 24 h twice. The resulting BSA-GlcFuc, BSA-Glc, and BSA-Fuc were concentrated to 1 mg/ml using Amicon ultra centrifugal filters 3000 MWCO.

### Testing of polyclonal anti-GlcFuc IgY

For the ELISA assays, BSA-GlcFuc, BSA-Fuc, and BSA were diluted at different concentrations in PBS supplemented with 1% BSA and 0.1% NaN_3_. Samples were incubated on ELISA plate at 4 °C overnight. Following blocking in PBST (PBS, pH 7.4, 0.05% Tween-20) with 10% BSA at RT for 1 h, the wells were incubated with 1 μg/ml purified anti-GlcFuc antibody in PBST + 10% BSA for 2 h. The secondary antibody (rabbit anti-chicken IgY HRP-conjugated) was diluted 1:100,000 in blocking buffer and incubated for 2 h at RT.

Signal was developed by using OPD (o-phenylenediamine dihydrochloride) reagent and read at the wavelength of 492 nm and 620 nm. For Western blot, 0.5 μg of protein per lane was loaded on a 8% SDS-PAGE gel with 4% stacking, transferred onto nitrocellulose membranes, and blocked in blocking buffer (% milk in PBST) for 1 h at RT. Anti-GlcFuc antibody was diluted in blocking buffer to 1 μg/ml and incubated for 1 h at RT. The secondary antibody (rabbit anti-chicken IgY HRP-conjugated) was diluted 1:5000 in blocking buffer and incubated for 1 h at RT. Blots were developed using SuperSignal West Pico (Pierce).

### Western blot

For total cell lysate, extracellular tachyzoites were harvested by centrifugation, washed twice in PBS and lysed in 1x reducing SDS-PAGE loading buffer with additional 0.1M DTT. Lysates were heated 10 min at 96 °C and about 5x10^6^ cells equivalent / lane were loaded on 8-16% TGX gels (Life Technologies). PAGE separated proteins were blotted on PVDF and the membranes were then blocked in 50 mM TrisHCl, 0.15 M NaCl, 0.25% BSA, 0.05% NP-40 pH 7.4 (61). ß-elimination on blot was performed as previously described (62). Briefly, after blotting the membrane was washed in PBS and incubated 16 h in 55 mM NaOH rotating at 40 °C. After rinsing in water, the membrane was blocked as described above. Both primary and secondary antibodies were diluted in blocking buffer as follow: mouse MAb anti-MIC2 6D10 1:5000, mouse MAb anti-*α*-tubulin 1:800 (DHSB), chicken anti-GlcFuc 1.1 μg/ml, anti-mouse HRP-conjugated (BioRad) 1:1000, and anti-chicken HRP 1:5000. Blots were developed by chemiluminescence (SuperSignal West Pico PLUS) using an ImageQuant LAS4000 imager (GE Healthcare). Quantification was performed using the ImageQuant TL software. The average of three biological repeats ± SD is shown. A two-tailed, heteroscedastic Student t test was used to calculate the *p* values (t.test function in Excel).

### Glycopeptide analysis by nUPLC-MS/MS (HCD)

Secreted proteins were isolated and prepared for nano Ultra Performance Liquid Chromatography Tandem Mass Spectrometry (nUPLC-MS/MS) analysis on a Thermo Q Exactive quadrupole Orbitrap MS as previously described (7, 25). Briefly, 4x10^10^ tachyzoites were isolated and resuspended in DMEM with 1% ethanol. After removal of parasites by centrifugation, the supernatant was filtered and concentrated. About 110 μg total protein was precipitated in 0.1 M ammonium acetate in MeOH for at least 16 h at -20 °C. After washing, the sample was dried by speed vacuum (Speed Vac Plus, Savant) and re-suspended in 50 mM ammonium bicarbonate, pH 8. The proteins were then reduced, alkylated with iodoacetamide and digested with trypsin and the resulting peptides were desalted on a silica C18 column (Nest Group). After speed vacuum drying, the sample was reconstituted in 99% Water/1% Acetonitrile (ACN)/0.1% Formic Acid (FA). For the initial discovery method, a two-μL aliquot was analyzed as previously described (25). Precursor ions containing the O-fucosylation peptide motif obtained from the Mascot search were analyzed by targeted nUPLC-MS/MS (HCD) in Parallel Reaction Monitoring (PRM) mode with an inclusion list containing both the *m/z* value and retention times of the ions of interest. The PRM experiment was set up as follows: a full mass scan was acquired with 70,000 resolution ^@^ *m/z* 200, AGC target 1 x 10^6^, 100 ms maximum injection time, 1 μscan/spectrum over the range *m/z* 370-2000. The selected precursor masses were calibrated with background ions at *m/z* 371.1012, 391.2843, 429.0887, and 445.1200 as the Lock Masses. The full mass range scan event was followed by seven PRM scan events, each with a different normalized collision energy. The PRM scan events were recorded with 17,500 resolution ^@^ *m/z* 200, AGC target 1 x 10^6^, 150 ms maximum injection time, 2 μscan/spectrum and their first mass was fit to *m/z* 100. The isolation window was set to 2 *m/z* units (Th) with an isolation offset of 0.4. An inclusion list with the *m/z* and retention times of the ions of interest was loaded, and the inclusion list feature was set to “on”. Ion dissociation/activation was achieved by quadrupole/beam-type CAD or HCD (Higher-Energy Collisional Dissociation). For each of the seven PRM scan events, a different normalized collision energy (NCE) was used. The NCE was set from 10 eV to 40 eV by increments of 5 eV.

### Glycopeptides analysis by nUPLC-MS/MS (ETD)

Precursor ions of most interest that were identified in the initial HCD experiment were used for a targeted nUPLC-MS/MS Electron Transfer Dissociation (ETD) experiment. An inclusion list of *m/z* with retention times was used in the ETD targeted experiment. Five-μL aliquots of the sample were injected into a nanoAcquity-UPLC (Waters) equipped with reversed phase columns: 5-μm Symmetry C18, 180 μm x 20 mm, trap column and 1.7 μm BEH130 C18, 150 μm x 100 mm, analytical column (Waters) and separated as described before (25). The nano-UPLC was connected online to an LTQ-Orbitrap XL Mass Spectrometer (Thermo Scientific) equipped with a Triversa NanoMate (Advion) electrospray ionization (ESI) source. The mass spectrometer was operated in the positive ion mode. The MS and tandem MS data were acquired in automatic Data Dependent “top 3” mode. The molecular ions were detected in the Orbitrap mass analyzer at a resolution of 30,000 ^@^ *m/z* 400, over the full scan range *m/z* 300-2000, 1 μscan/spectrum, maximum injection time (ion accumulation time) of 500 ms with a target automatic gain control (AGC) of 500.0 ion population. The selected precursor masses were calibrated with background ions at *m/z* 371.1012, 391.2842 and 445.1200 as the Lock Masses. The top three most abundant ions with charge state higher than 2 in the survey MS scan were selected, using a 2-Da isolation window, and were fragmented by ETD in the linear ion trap (LTQ) mass analyzer. The fluoranthene radical anions were accumulated for a maximum time of 125 ms and AGC of 500,000 prior to the ETD reaction. The ETD activation time was 125 ms, with supplemental activation of 15 V. For *m/z* 737.3218 (2+) the fragments were detected in the Orbitrap mass analyzer with 2 μscan/spectrum, maximum injection time of 700 ms with AGC of 200.0 ion population. For *m/z* 586.7708 (2+) the fragments were detected in the LTQ mass analyzer with 3 μscan/spectrum, maximum injection time of 150 ms with AGC of 10,000. All mass spectrometry data have been deposited to the ProteomeXchange Consortium (<u>http://proteomecentral.proteomexchange.org</u>) via the PRIDE partner repository with the dataset identifier PXD010622 (63).

### Plaque assay

Host cell monolayers on 6-well plates were infected with 250 parasites/well of either parental or knockout strains. Parasites were allowed to grow at 37 °C, 5% CO_2_ for 8 days. After washing in PBS, cells were fixed in ice-cold methanol at -20 °C for at least 20 min and stained with Giemsa for 10 min. After the cells were washed with ddH2O and air-dried, plates were imaged on an ImageQuant LAS400. Plaque areas were determined using the ImageQuant TL software. Plaque numbers shown are the average of four biological repeats ± SD. A two-tailed, homoscedastic Student t test was used to calculate the *p* values (t.test function in Excel).

### Attachment/invasion assay

The protocol was adapted from (46). Briefly, HFF monolayers on coverslip (24-well plate) were infected with 5x10^6^ tachyzoites/ well from either parental or knockout strains. The plate was centrifuged 5 min at 200 x *g* and incubated 30 min at 37 °C. After washing with PBS to remove unattached parasites, the coverslips were fixed in 4% PFA in PB for 20 min on ice and blocked in 3% BSA/ PBS for 1 h at RT. Extracellular (attached) parasites were labeled with MAb anti-SAG1 T4 1E5 1:300 (BEI Resources). Cells were then permeabilized (10 min, 0.25% TX100/PBS) and blocked once more (3% BSA/PBS, 30 min at RT). All parasites (attached and invaded) were labeled with rabbit anti-ß-tubulin 1:1000 (this step was required as the parental strain does not express mGFP like the knockout strains analyzed in this study). Anti-mouse AlexaFluor594 and anti-rabbit AlexaFluor488 were used as secondary antibodies. Coverslips were mounted on glass slides using Prolong Diamond with DAPI (Molecular Probes) and examined on a Zeiss AXIO inverted microscope. Six randomly selected fields were imaged with a total magnification of 400x. Attached (red + green) and invaded (green) parasites were counted. Parasite counts were normalized to the number of host cell nuclei in each field and the data from three biological repeats are shown. A two-tailed, heteroscedastic Student t test was used to calculate the *p* values (t.test function in Excel).

### Egress assay

The protocol was adapted from (13) and (45). Briefly, HFF monolayers on coverslip (24-well plate) were infected with 1x10^5^ tachyzoites/ well from either parental or knockout strains and parasites were allowed to grow for 24-28h. To induce egress, cells were incubated for 5 min at 37°C with 2 μM A23178 (Sigma) in pre-warmed DMEM (no FBS). Coverslips were immediately fixed in 4% PFA in PB for 20 min on ice and blocked in 3% BSA/ PBS for 1 h at RT. Labeling and imaging were performed as described for the invasion/attachment assay with the difference that 8-10 fields were scored in each biological repeat. The average of three independent experiments ± SD is shown. A two-tailed, heteroscedastic Student t test was used to calculate the *p* values (t.test function in Excel).

## Acknowledgements

We would like to thanks Prof. Maryse Lebrun for the AMA4 antibody and Prof. Markus Meissner, Prof. Vern Carruthers, and Prof. Marc Jan Gubbels for reagents and helpful discussions.

## Author contributions

J.S. and G.B. conceived the study and designed the experiments. D.R.L. performed the mass spectrometry analysis; C.M.H. isolated the secreted proteins; Y.Z., L.K.M., and M.J.S. produced and characterized the antibody. G.B. performed all remaining experiments. G.B., J.S., D.R.L., C.A.N., F.H.R., and C.E.C. performed the data analysis, and all authors contributed to the writing of the manuscript.

## The abbreviations used are

MIC2: microneme protein 2
TSR: thrombospondin repeat
POFUT2: protein O-fucosyltransferase 2
NST2: nucleotide sugar transporter 2
M2AP: MIC2 associated protein
mGFP: green fluorescent protein
mRFP: red fluorescent protein
AAL: Aleuria aurantia lectin
KLH: keyhole limpet hemocyanin
dHex: deoxyhexose
Hex: hexose
Fuc: fucose
Glc: glucose
Man: mannose
HCD: higher collision energy dissociation
ETD: electron transfer dissociation
PV: parasitophorous vacuole

